# In the Absence of Apoptosis, Myeloid Cells Arrest When Deprived of Growth Factor, But Remain Viable by Consuming Extracellular Glucose

**DOI:** 10.1101/198754

**Authors:** Li Dong, Boris Reljic, Jen G. Cheung, Elizabeth S. Ng, Lisa M. Lindqvist, Andrew G. Elefanty, David L. Vaux, Hoanh Tran

**Author notes:** Address correspondence to Hoanh Tran.

## Abstract

Withdrawal of the growth factor interleukin 3 from IL3-dependent myeloid cells causes them to undergo Bax/Bak1-dependent apoptosis, whereas factor-deprived *Bax*^-/-^*Bak1*^-/-^ cells remain viable, but arrest and shrink. It was reported that withdrawal of IL3 from *Bax*^-/-^*Bak1*^-/-^ cells caused decreased expression of the glucose transporter Glut1, leading to reduced glucose uptake, so that arrested cells required Atg5-dependent autophagy for long-term survival. In other cell types, a decrease in Glut1 is mediated by the thioredoxin-interacting protein Txnip, which is induced in IL3-dependent myeloid cells when growth factor is removed. We mutated *Atg5* and *Txnip* by CRISPR/Cas9 and found that Atg5-dependent autophagy was not necessary for the long-term viability of cycling or arrested *Bax*^-/-^*Bak1*^-/-^ cells, and that Txnip was not required for the decrease in Glut1 expression in response to IL3 withdrawal. Surprisingly, Atg5-deficient *Bax/Bak1* double mutant cells survived for several weeks in medium supplemented with 10% fetal bovine serum (FBS), without high concentrations of added glucose or glutamine. When serum was withdrawn, the provision of an equivalent amount of glucose present in 10% FBS (~0.5 mM) was sufficient to support cell survival for more than a week, in the presence or absence of IL3. Thus, *Bax*^-/-^*Bak1*^-/-^ myeloid cells deprived of growth factor consume extracellular glucose to maintain long-term viability, without a requirement for Atg5-dependent autophagy.

## Introduction

Mammalian cells import a variety of extracellular nutrients to support their survival, growth and proliferation. *In vitro*, cells are typically propagated in medium supplemented with glucose, glutamine and fetal bovine serum. To generate chemical energy, glucose can be metabolized by oxidative phosphorylation in the mitochondria or by anaerobic glycolysis. Glutamine can be used as an energy source through glutaminolysis, and can also be used, together with a multitude of factors present in serum, to allow the synthesis of essential macromolecules for cell division.

Factor dependent myeloid (FDM) progenitor cell lines^1, 2^ require the hematopoietic growth factor interleukin 3 (IL3) to survive and multiply in culture. When IL3 is not provided, FDM cells undergo Bax/Bak1-dependent apoptosis. FDM cells derived from *Bax*^-/-^*Bak1*^-/-^ mice survive in the absence of IL3, but they undergo cell cycle arrest, similar to those over-expressing Bcl-2^3, 4^. It has been reported that to survive longer than 4 days, IL3-deprived *Bax*^-/-^*Bak1*^-/-^ cells required Atg5-dependent autophagy^4^, an intracellular catabolic process that can support the survival of starved or arrested cells^5^.

Cells import glucose when it is available and growth factors can stimulate the amount of glucose that is consumed. *Bax*^-/-^*Bak1*^-/-^ FDM cells arrested by IL3 withdrawal decrease expression of the glucose transporter Glut1, which was proposed to reduce glucose uptake and metabolism^4^. Several mechanisms have been put forward to explain the regulation of expression and plasma membrane translocation of Glut1. Activation of autophagy was shown to be required for the efficient surface expression of Glut1 in mouse embryonic fibroblasts cultured under hypoxic conditions^6^. In human hepatocytes subjected to glucose starvation, a mechanism involving proteasomal degradation of the thioredoxin-interacting protein Txnip was required for Glut1 stabilization^7^. Moreover, it was shown that Txnip directly binds to Glut1 and promotes its internalization, thus reducing Glut1 on the plasma membrane and the ability of cells to take up glucose^7^.

In this study, we employed *Bax*^-/-^*Bak1*^-/-^ FDM cells to genetically test whether Atg5-dependent autophagy is required for the sustained survival of arrested cells deprived of growth factor. We also asked whether autophagy and/or Txnip are required for the reduction in Glut1 expression and glucose uptake by arrested myeloid cells.

## Results

### Bax^-/-^Bak1^-/-^ myeloid cells shrink and arrest when deprived of IL3, but survive

Factor dependent myeloid (FDM) cells require interleukin 3 (IL3) to survive and proliferate^2, 4, 8^. When deprived of IL3 for 4 days, >80% of wild-type *Bax*^+/+^*Bak1*^+/+^ FDM cells died, as indicated by loss of plasma membrane integrity and uptake of propidium iodide (Figs. 1A and Supplementary S1A). By contrast, and consistent with the findings of Lum *et al*.^4^, *Bax*^-/-^*Bak1*^-/-^ FDM cells survived in the absence of IL3 for more than 14 days. After 48 h IL3 withdrawal, these cells shrank in size and arrested (Fig. 1, B and C). This quiescent state was reversible, because when IL3 was returned, the cells increased in size and resumed cycling. Thus, in the absence of Bax/Bak1-mediated apoptosis, FDM cells can survive for weeks without IL3, but they shrink and arrest.

**Figure 1.**
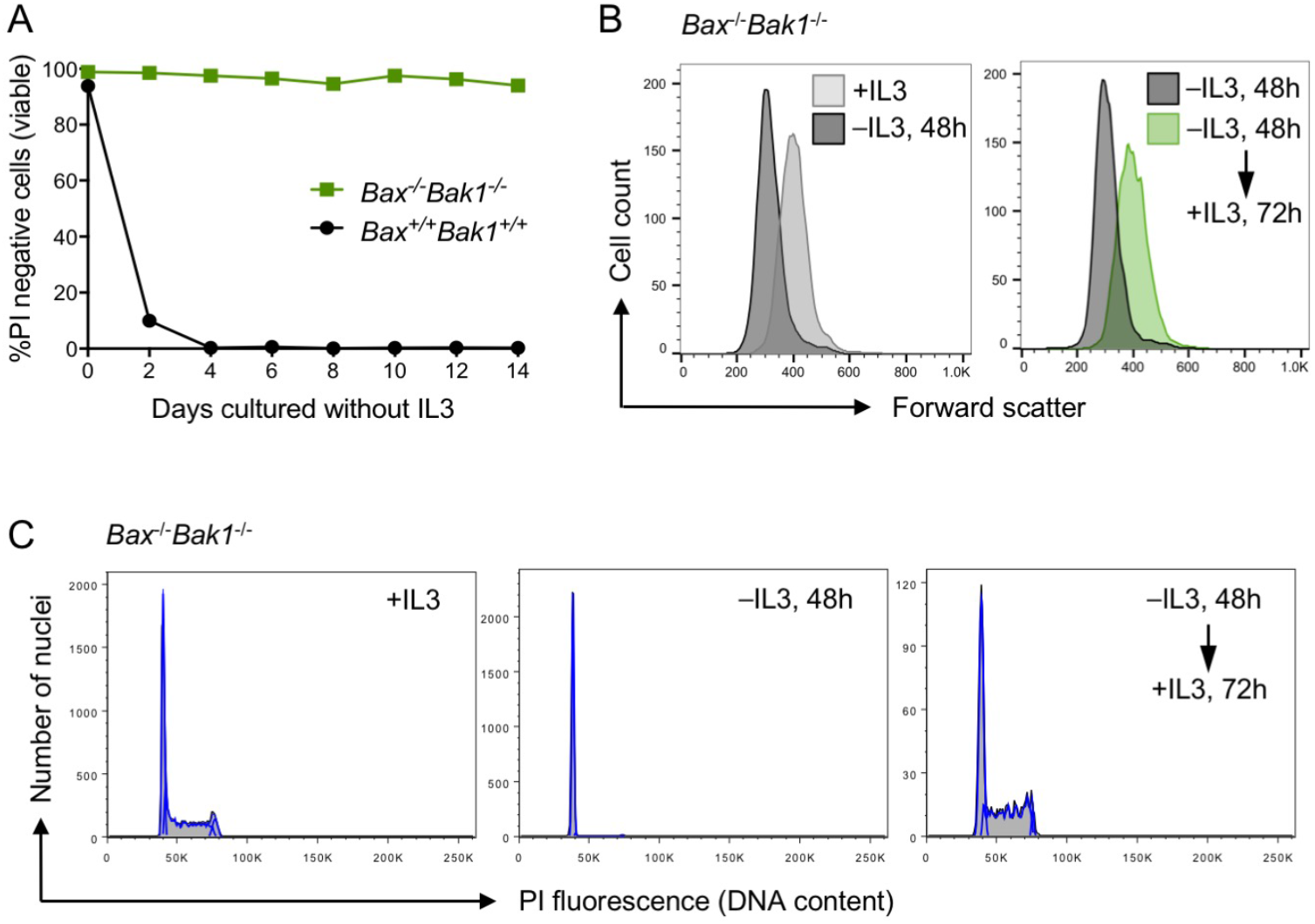
*Bax*^-/-^*Bak1*^-/-^ FDM cells shrink and arrest but remain viable when cultured without IL3. **A.** *Bax*^-/-^*Bak1*^/-^ FDM cells survive long-term in the absence of IL3. *Bax*^+/+^*Bak1*^+/+^ wild-type (black circles) and *Bax^-/-^Bak1^-/-^* (green squares) FDM cells were cultured without IL3. At the indicated times, cells were harvested, resuspended in PBS containing propidium iodide (PI) and analyzed by flow cytometry. **B.** IL3 withdrawal caused cells to shrink. PI negative populations of *Bax*^-/-^*Bak1*^-/-^ cells cultured for 48 h in the presence (light gray) or absence (dark gray) of IL3, and the latter that had IL3 returned for 72 h (green), were compared by forward scatter as an indicator of cell size. C. IL3 withdrawal caused cell cycle arrest, which was reversed by re-addition of IL3. Nuclei from *Bax*^-/-^*Bak1*^-/-^ cells cultured as described in (B) were stained with PI to indicate DNA content by flow cytometry.

### Bax^-/-^Bak1^-/-^ myeloid cells arrested by deprivation of IL3 do not require Atg5-dependent autophagy to survive

During macroautophagy, LC3B-I is lipidated by a multiprotein complex containing Atg5 to form LC3B-II, which is incorporated into the growing autophagosome, and is commonly used as a marker of autophagy^9^. Lysates of *Bax*^-/-^*Bak1*^-/-^ FDM cells, whether they were cultured in the presence or absence of IL3, contained similar amounts LC3B-II, suggesting that autophagy is occurring in both cycling and arrested cells (Fig. 2A; lanes 1, 3 and 4). Notably, the low levels of both LC3B isoforms I and II in *Bax*^-/-^*Bak1*^-/-^ cells cultured in serum, amino acid and glucose-free Hank’s balanced salt solution (HBSS; Fig. 2A, lane 2) is indicative of a strong increase in autophagic flux, which causes the coincident degradation of LC3B proteins after autophagosomes fuse with lysosomes^9^.

**Figure 2.**
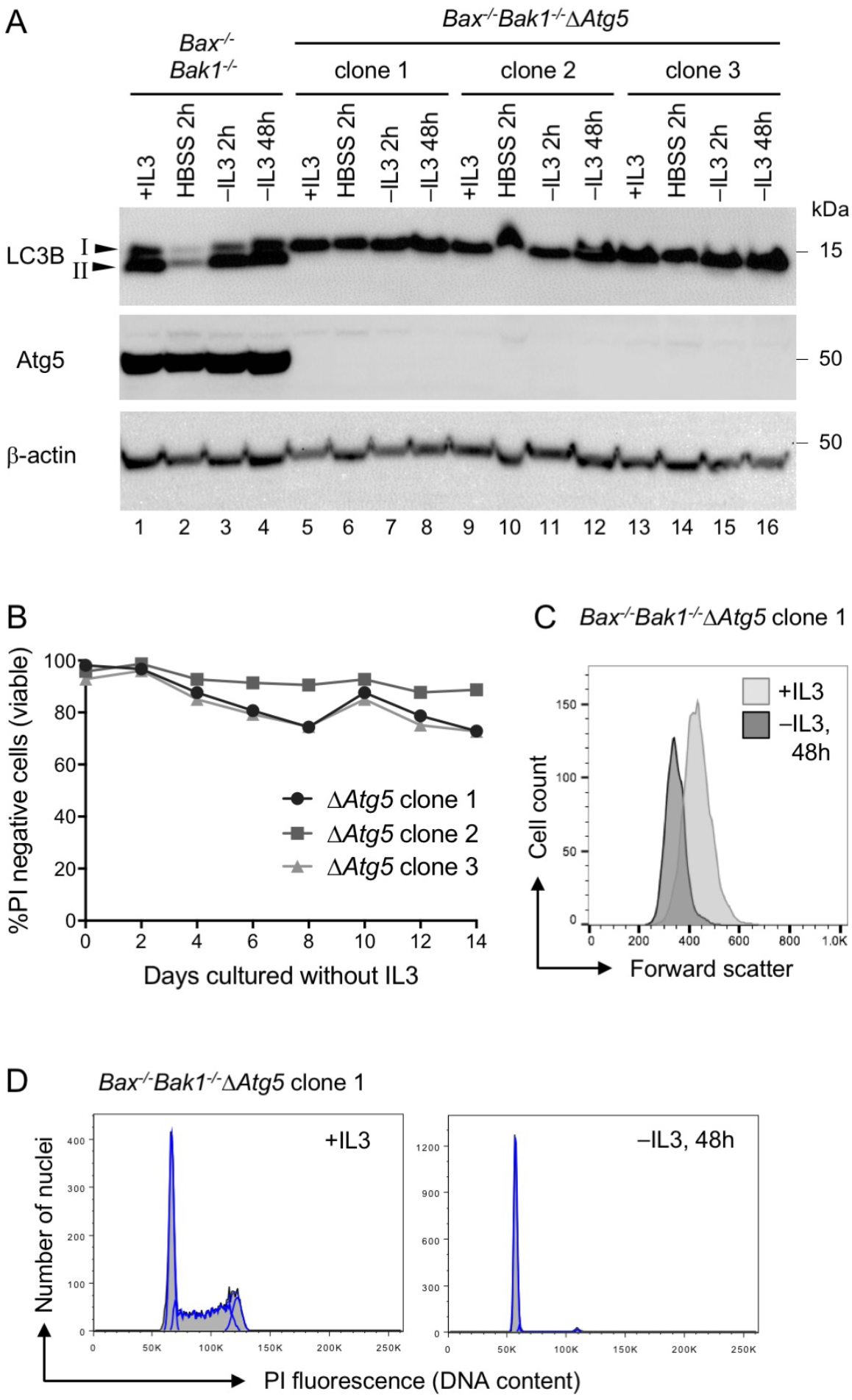
*Bax*^-/-^*Bak1*^-/-^Δ*Atg5* FDM cells shrink and arrest, but remain viable when IL3 is withdrawn. **A.** Whole cell lysates from *Bax*^-/-^*Bak1*^-/-^ cells and 3 *Bax*^-/-^*Bak1*^-/-^Δ*Atg5* clonal lines cultured in the presence of IL3, in Hank’s Buffered Saline Solution (HBSS) for 2 h, or in the absence of IL3 for 2 or 48 h, were subjected to Western blotting. The Atg5 immunoblot demonstrates successful CRISPR/Cas9-mediated deletion of *Atg5*. The LC3B immunoblot demonstrates lack of LC3B lipidation (isoform II) and autophagic flux (HBSS treatment) in Δ*Atg5* cell clones. **B.** *Bax*^-/-^*Bak1*^-/-^Δ*Atg5* FDM cells survive long-term in the absence of IL3. The indicated *Bax*^-/-^*Bak1*^-/-^Δ*Atg5* clones were cultured without IL3 and viability was determined by PI exclusion and flow cytometry at the indicated times. **C.** *Bax*^-/-^*Bak1*^/-^Δ*Atg5* cells (clone 1) were cultured for 48 h in the presence or absence of IL3 and PI negative populations were compared by forward scatter to indicate differences in cell size. **D.** Cells from *Bax*^-/-^*Bak1*^-/-^Δ*Atg5* clone 1 were cultured for 48 h in the presence or absence of IL3 then processed in PI buffer to measure DNA content by flow cytometry.

To determine if Atg5-dependent autophagy was needed to support the sustained survival of IL3-starved cells, we used CRISPR/Cas9 to delete *Atg5* in *Bax*^-/-^*Bak1*^-/-^ cells, generating *Bax*^-/-^*Bak1*^-/-^Δ*Atg5* triple knockout FDM cell clones. Western blotting confirmed in three *Bax*^-/-^*Bak1*^-/-^Δ*Atg5* clones the absence of the LC3B-II isoform, whether they were cultured with or without IL3, or in HBSS (Fig. 2A). Moreover, levels of LC3B-I in *Bax*^-/-^*Bak1*^-/-^Δ*Atg5* clones remained similar across the different culture conditions. These results indicate that Atg5-dependent autophagy and autophagic flux are blocked in the *Bax*^-/-^*Bak1*^-/-^Δ*Atg5* lines. After 14 days cultured without IL3, all *Bax*^-/-^*Bak1*^-/-^Δ*Atg5* clones remained ~80% viable (Fig. 2B). Similar to *Bax*^-/-^*Bak1*^-/-^ FDMs, *Bax*^-/-^*Bak1*^-/-^Δ*Atg5* cells decreased in size and arrested within 48 h of growth factor withdrawal (Fig. 2, C and D).

These results confirm that IL3 receptor signals are needed for *Bax*^-/-^*Bak1*^-/-^ FDM cells to maintain their normal size and divide, but show that Atg5-dependent autophagy is not required for their survival, whether they are cycling or are arrested by IL3 withdrawal. Furthermore, because culturing without IL3 caused a similar decrease in size of the arrested *Bax*^-/-^*Bak1*^-/-^ to that of the *Bax*^-/-^*Bak1*^-/-^Δ*Atg5* cells, this shrinkage is not due to Atg5-dependent autophagy.

### Bax^-/-^Bak1^-/-^ myeloid cells arrested by IL3 deprivation downregulate Glut1 independently of Txnip or Atg5 and survive in the absence of added glucose

Lum *et al*. reported that when IL3 growth factor was removed, *Bax*^-/-^*Bak1*^-/-^ FDM cells decreased expression of the glucose transporter Glut1^4^. A possible mechanism for this was proposed by Wu *et al*., who found that thioredoxin-interacting protein (Txnip) could not only bind to Glut1 and induce its internalization through clathrin-coated pits, thus removing it from the plasma membrane, but could also reduce the level of Glut1 messenger mRNA^7^. Consistent with Lum *et al*. and Wu *et al*., levels of Txnip increased, and those of Glut1 decreased, in lysates of *Bax*^-/-^*Bak1*^-/-^ cells 48 h following removal of IL3 (Fig. 3A, lane 2).

**Figure 3.**
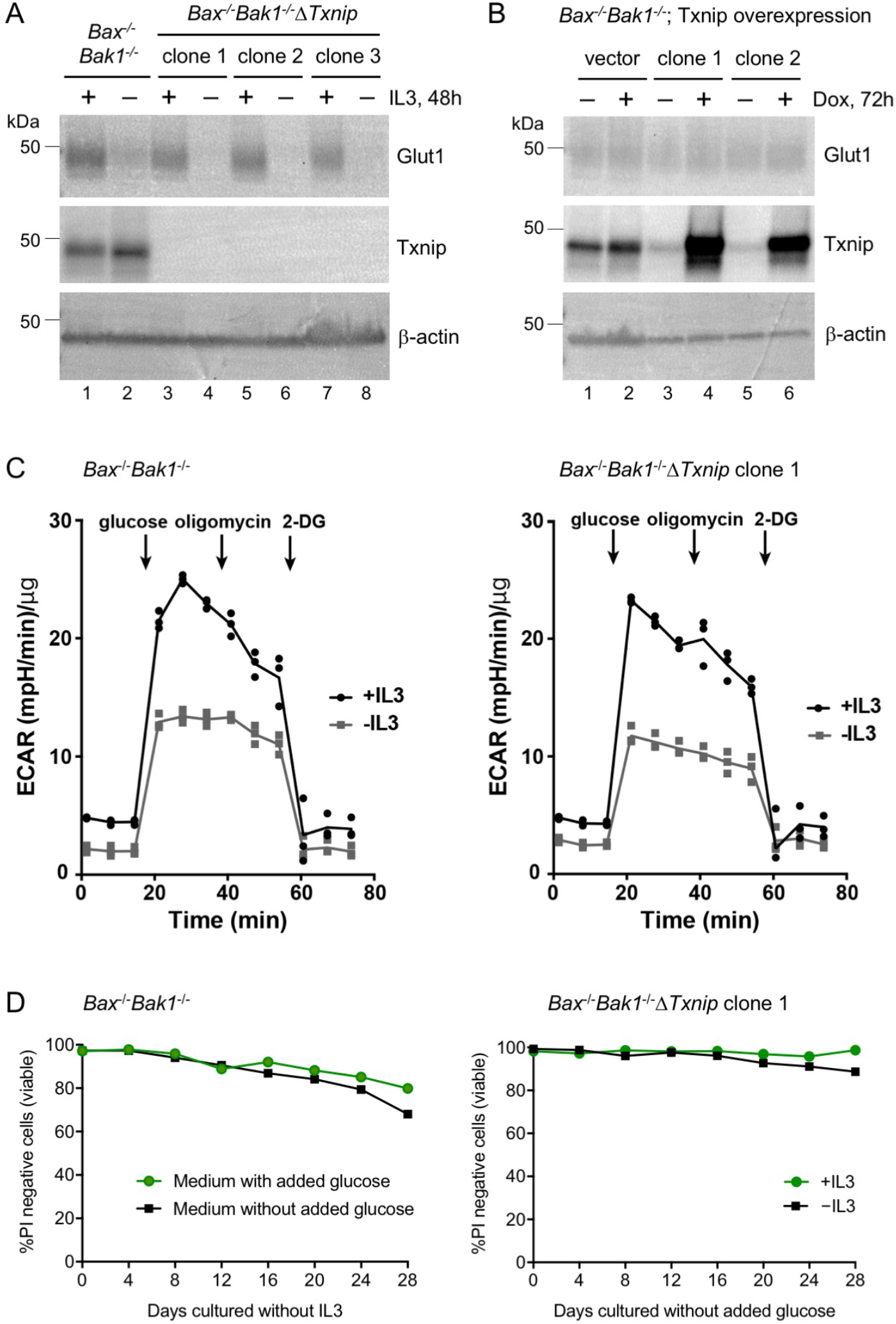
*Bax*^-/-^*Bak1*^/-^ FDM cells arrested by IL3 deprivation survive in the absence of added glucose. **A.** Txnip is not required for the decrease in Glut1 following IL3 withdrawal. Whole cell lysates from *Bax*^-/-^*Bak1*^-/-^ and 3 independent *Bax*^-/-^*Bak1*^-/-^Δ*Txnip* FDM cell clones cultured for 48 h in the presence or absence of IL3 were analyzed by Western blotting to detect Glut1 and Txnip protein. **B.** Forced expression of Txnip is not sufficient to decrease Glut1 protein levels. Independent *Bax*^-/-^*Bak1*^-/-^ FDM clones bearing a doxycycline (dox)-inducible FLAG-Txnip construct or empty vector control were cultured for 72 h in the absence or presence of dox. Whole cell lysates were prepared and analyzed by Western blotting to detect Glut1 and Txnip protein. **C.** *Bax*^-/-^*Bak1*^-/-^ and *Bax*^-/-^*Bak1*^-/-^Δ*Txnip* cells can take up and metabolise glucose. Cells cultured for 72 h in the presence or absence of IL3 were analyzed on the Seahorse bioanalyzer to measure the extracellular acidification rate (ECAR) as an indicator of glucose metabolism. The increased ECAR following addition of glucose signifies increased glycolytic flux in cells. Oligomycin inhibits mitochondrial ATP synthase and the change in ECAR upon its addition represents the glycolytic reserve capacity of cells. Glycolysis is inhibited by 2-deoxyglucose (2-DG) so upon its addition, the ECAR values are produced from non-glycolytic sources. Equal numbers of cells were seeded for analysis. To correct for the reduced size of IL3-starved cells, protein concentration was assayed and the ECAR values were normalized to 1 μg total protein per sample. Curves represent the mean of 3 replicate samples depicted as solid symbols for each time point. **D.** *Bax*^-/-^*Bak1*^-/-^ and *Bax*^-/-^*Bak1*^-/-^Δ*Txnip* cells deprived of IL3 survive long-term in the absence of added glucose. Cells were cultured without or with IL3 in medium supplemented with 10% FBS, 4 mM glutamine and 5.5 mM glucose (medium with added glucose) or in medium supplemented with 10% FBS and 4 mM glutamine (medium without added glucose). Cell viability was monitored by PI exclusion and flow cytometry at the indicated times.

To determine whether expression of Txnip caused or correlated with changes in Glut1 levels, we deleted *Txnip* in *Bax*^-/-^*Bak1*^-/-^ FDM cells using CRISPR/Cas9. Western blotting confirmed the absence of Txnip protein in three *Bax*^-/-^*Bak1*^-/-^Δ*Txnip* clonal lines (Fig. 3A). We found that Glut1 levels were unaffected by deletion of *Txnip*, whether IL3 was added or not, indicating that the increase in Txnip was not required for the decrease in Glut1 when cells were cultured without IL3. Furthermore, similar to *Bax*^-/-^*Bak1*^-/-^ cells, withdrawal of IL3 caused G1 arrest in *Bax*^-/-^*Bak1*^-/-^Δ*Txnip* cells within 48 h (Supplementary Fig. S1A). Because induced over-expression of FLAG-Txnip in *Bax*^-/-^*Bak1*^-/-^ cells had no impact on levels of Glut1 protein or cell cycle progression (Figs. 3B and Supplementary S1B), an increase in Txnip is not sufficient to reduce Glut1. Therefore, in FDM cells, the abundance of Txnip has no effect on levels of Glut1, whether they are treated with IL3 or not. Of note, IL3 deprivation caused specific downregulation of Glut1, but not Glut4, protein in *Bax*^-/-^*Bak1*^-/-^ and *Bax*^-/-^*Bak1*^-/-^Δ*Atg5* clones; and levels of Glut1 were similar among proliferating Bax/Bak1-null cells of all genotypes (Figs. 3A, B and Supplementary Fig. S1C). Together, these results indicate that although IL3 can promote the expression of Glut1, neither Txnip nor Atg5-dependent autophagy is required for the decrease in Glut1 when growth factor is removed from myeloid cells, and neither absence nor overexpression of Txnip affects the levels of Glut1.

Because Glut1 levels are reduced in IL3-starved *Bax*^-/-^*Bak1*^-/-^ FDM cells, we asked whether glucose utilization was reduced. Proliferating *Bax*^-/-^*Bak1*^-/-^ cells in the presence of IL3 responded strongly to the addition of glucose, as indicated by the increased extracellular acidification rate (ECAR, Fig. 3C), signifying that they readily consumed and metabolized the added glucose. Following 3 days cultured without IL3, the rate of glucose metabolism in *Bax*^-/-^*Bak1*^-/-^ cells was reduced, but they nevertheless still responded to the addition of glucose. With or without IL3, the rates of glucose metabolism in *Bax*^-/-^*Bak1*^-/-^ lines that also lacked Txnip or Atg5 were similar to the rates of the parental Bax/Bak1-null cells (Figs. 3C and Supplementary S1D). When cells were treated with oligomycin to block oxidative phosphorylation shortly after they reached their maximal glycolytic rate, the steady decline in ECAR continued relatively unchanged until glycolysis was inhibited with 2-deoxyglucose (2-DG), causing ECAR values to fall rapidly to baseline. These results indicate that FDM cells rely predominantly on glycolysis for energy production, and when deprived of IL3, they retain the capacity to import and metabolise glucose, albeit at a reduced level.

Because IL3-starved *Bax*^-/-^*Bak1*^-/-^ FDMs took up and metabolized glucose, we hypothesised that this was sufficient to support their sustained survival despite the decrease in Glut1. For FDM cell culture, we used DMEM medium supplemented with 4 mM glutamine, 5.5 mM glucose and 10% fetal bovine serum (FBS), estimated to contain ~0.5 mM glucose (Supplementary Fig. S1E). To test their requirement for glucose, the cells were cultured in medium supplemented with 4 mM glutamine and 10% FBS, but lacking the added 5.5 mM glucose. Surprisingly, > 80% of IL3-starved *Bax*^-/-^*Bak1*^-/-^ and *Bax*^-/-^*Bak1*^-/-^Δ*Txnip* cells remained viable for 3 weeks in culture without added glucose (Fig. 3D). Furthermore, when IL3 was present, these cells were able to proliferate (Supplementary Fig. S1F). These results suggest that FDM cells do not require high concentrations of glucose to maintain long-term viability, whether IL3 is present or not, and that they consume the glucose (and other nutrients) present in FBS and/or the added glutamine to support survival or to multiply when stimulated with growth factor.

### Bax^-/-^Bak1^-/-^ myeloid cells require glutamine to proliferate but can survive in its absence for weeks

When glucose is limiting, lymphoid cells metabolise glutamine to survive and proliferate^10^. We therefore asked whether IL3-dependent myeloid cells exhibited similar requirements for glutamine. In medium supplemented with 10% FBS but without added glucose or glutamine, with or without IL3, >95% of wild-type *Bax*^+/+^*Bak1*^+/+^ FDM cells died as indicated by uptake of PI by day 4 (Supplementary Fig. S2A), and consistent with their death being due to apoptosis, the nuclei of the cells had sub-2n DNA (Fig. 4A). Apoptosis was Bax/Bak1-dependent, because Bax/Bak1-null cells, including clones also lacking Atg5, did not die when cultured in medium without added glucose and glutamine, but they arrested instead, even when IL3 was provided (Fig. 4A). When cultured with neither added glucose nor glutamine, in the presence or absence of IL3, >80% of the arrested *Bax*^-/-^*Bak1*^-/-^ and *Bax*^-/-^*Bak1*^-/-^Δ*Atg5* cells survived for more than 12 days (Figs. 4B and Supplementary S2B).

**Figure 4.**
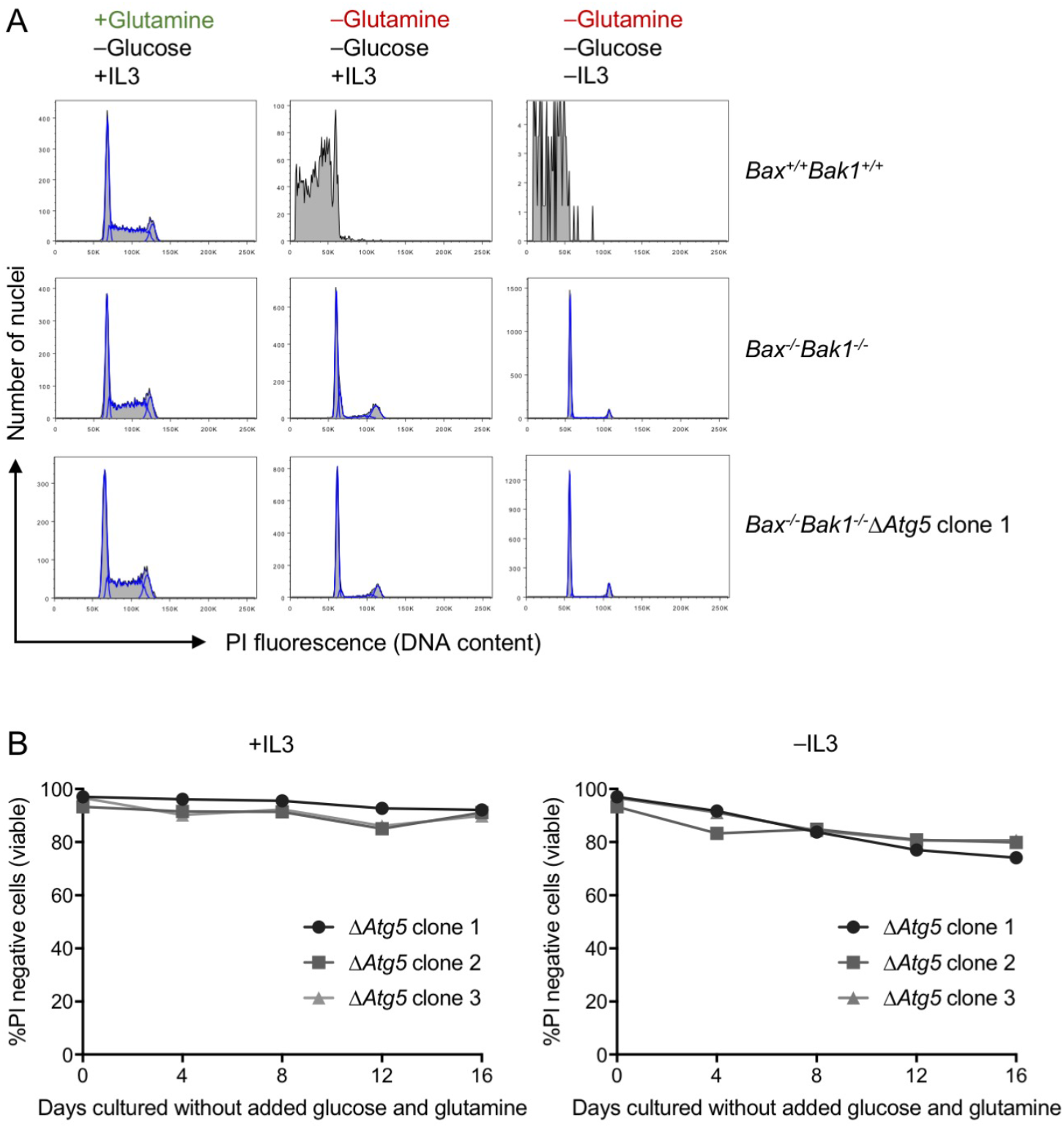
When starved of glutamine, *Bax*^+/+^*Bak1*^+/+^ FDM cells undergo apoptosis, whereas *Bax*^-/-^*Bak1*^-/-^ cells arrest, but can survive for weeks. **A.** *Bax*^+/+^*Bak1*^+/+^ cells undergo apoptosis in the absence of glutamine (upper panels), whereas *Bax*^-/-^*Bak1*^/-^ and *Bax*^-/-^*Bak1*^-/-^Δ*Atg5* cells survive in the absence of glutamine, but arrest even when IL3 is present (lower panels). Cells were cultured in medium supplemented with 10% FBS, with or without IL3 and 4 mM glutamine, in the absence of added glucose, for 4 days. Cells were processed in PI buffer to assess DNA content by flow cytometry. Note the absence of cells in S phase when Bax/Bak1-null cells were cultured without added glutamine. **B.** *Bax*^-/-^*Bak1*^-/-^Δ*Atg5* cells survive for weeks in the absence of glutamine. Three independent *Bax*^-/-^*Bak1*^-/-^Δ*Atg5* clones were cultured with or without IL3, in medium supplemented with 10% FBS, without added glucose or glutamine. At the indicated times, cells were assayed for viability by PI exclusion and flow cytometry.

When *Bax*^-/-^*Bak1*^-/-^ cells were starved of glucose and glutamine, most arrested in G1 with 2n DNA, but some arrested in G_2_/M with 4n DNA (Fig. 4A). When glutamine was added back to *Bax*^-/-^*Bak1*^-/-^ cells that had been cultured for 7 days in glucose- and glutamine-free medium, the arrested cells resumed cycling within 48 h in the presence of IL3, whereas adding back glucose had no effect (Supplementary Fig. S2C). These results indicate that IL3-treated *Bax*^-/-^*Bak1*^-/-^ myeloid cells require glutamine to proliferate, but can survive for weeks in its absence. Because *Bax*^-/-^*Bak1*^-/-^Δ*Atg5* cells behaved similarly to *Bax*^-/-^*Bak1*^-/-^ cells, Atg5-dependent autophagy is not required for the sustained survival of growth factor-deprived *Bax*^-/-^*Bak1*^-/-^ FDMs, even in cultures without high levels of glucose and glutamine.

### Bax^-/-^Bak1^-/-^ myeloid cells arrested by glutamine deprivation shrink in a manner independent of Atg5-mediated autophagy

Because *Bax*^-/-^*Bak1*^-/-^ and *Bax*^-/-^*Bak1*^-/-^Δ*Atg5* cells arrested by IL3 withdrawal decreased in size (Figs. 1B and 2C), we asked whether cells arrested by glutamine withdrawal also shrink. In medium containing 10% FBS but lacking added glucose and glutamine, IL3-treated *Bax*^-/-^*Bak1*^-/-^ and *Bax*^-/-^*Bak1*^-/-^Δ*Atg5* cells showed a gradual decrease in size to almost 1/3 of their original volume over a 20-day period (Fig. 5). Without IL3, the shrinkage was more acute and cells reached ~1/3 of their original volume within 7 days. These results suggest that IL3 provided a signal to *Bax*^-/-^*Bak1*^-/-^ myeloid cells arrested by glutamine deprivation to maintain a large size, and confirm that the shrinkage of these cells upon IL3 and/or nutrient withdrawal do not require Atg5-dependent autophagy.

**Figure 5.**
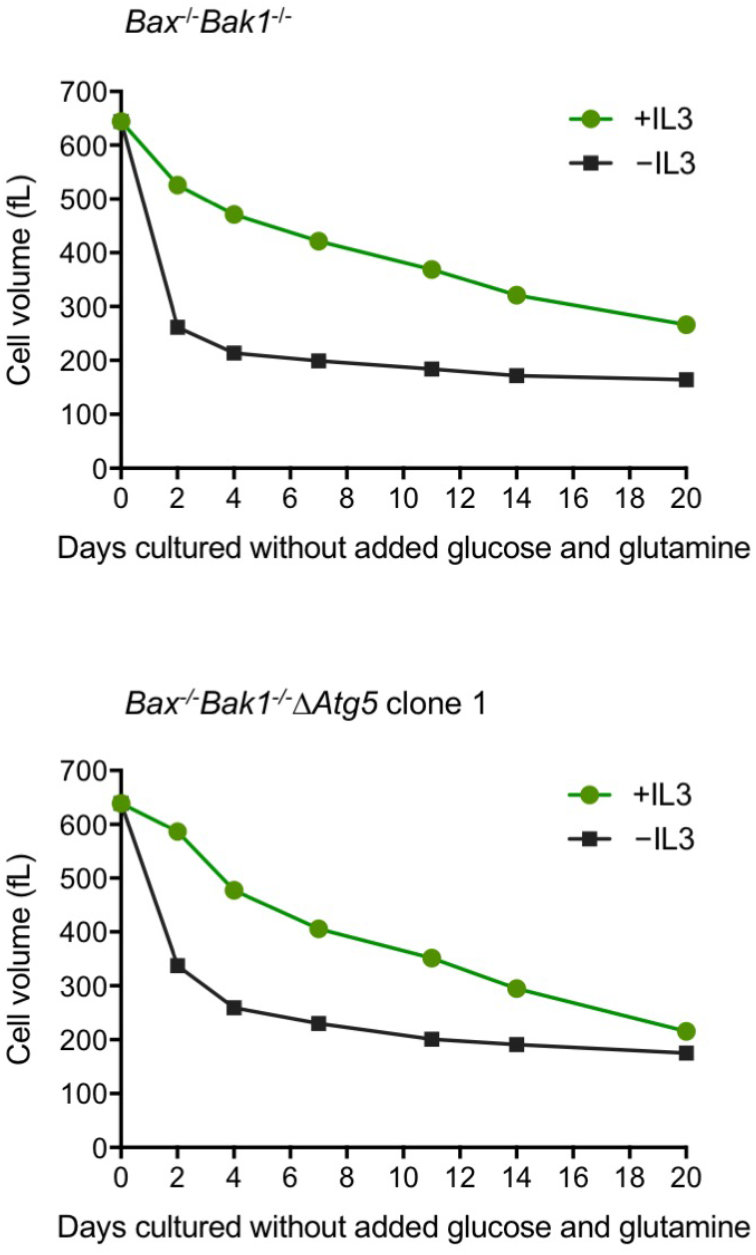
Withdrawal of IL3 or glutamine caused cell shrinkage independent of Atg5-mediated autophagy. *Bax*^-/-^*Bak1*^-/-^ and *Bax*^-/-^*Bak1*^-/-^Δ*Atg5* cells were cultured in medium supplemented with 10% FBS, with or without IL3, in the absence of added glucose and glutamine. At the indicated times, cell volume was measured in a Coulter-based CASY cell counter.

### Withdrawal of IL3 or glutamine caused reduced expression of Mcl-1 and key regulators of cell cycle progression

To elucidate how growth factor and nutrient deprivation impacted FDM cell survival and proliferation, we performed Western blotting to determine protein levels of apoptotic and cell cycle regulators. We found that levels of the pro-survival protein Mcl-1 were markedly reduced in *Bax*^-/-^*Bak1*^-/-^ and *Bax*^-/-^*Bak1*^-/-^Δ*Atg5* cells after 3 days in culture without added glucose and glutamine (Fig. 6A), suggesting that the decrease in normal levels of Mcl-1 allowed activation of Bax/Bak1 and the apoptosis of wild-type FDM cells observed in Fig. 4A. Consistent with this notion, addition of a specific Mcl-1 inhibitor S63845^11^ to *Bax*^+/+^*Bak1*^+/+^ cells was sufficient to cause significant apoptosis within 24 h, even when the cells were cultured with IL3 in complete medium containing 10% FBS and high concentrations of glucose and glutamine (Fig. 6B).

**Figure 6.**
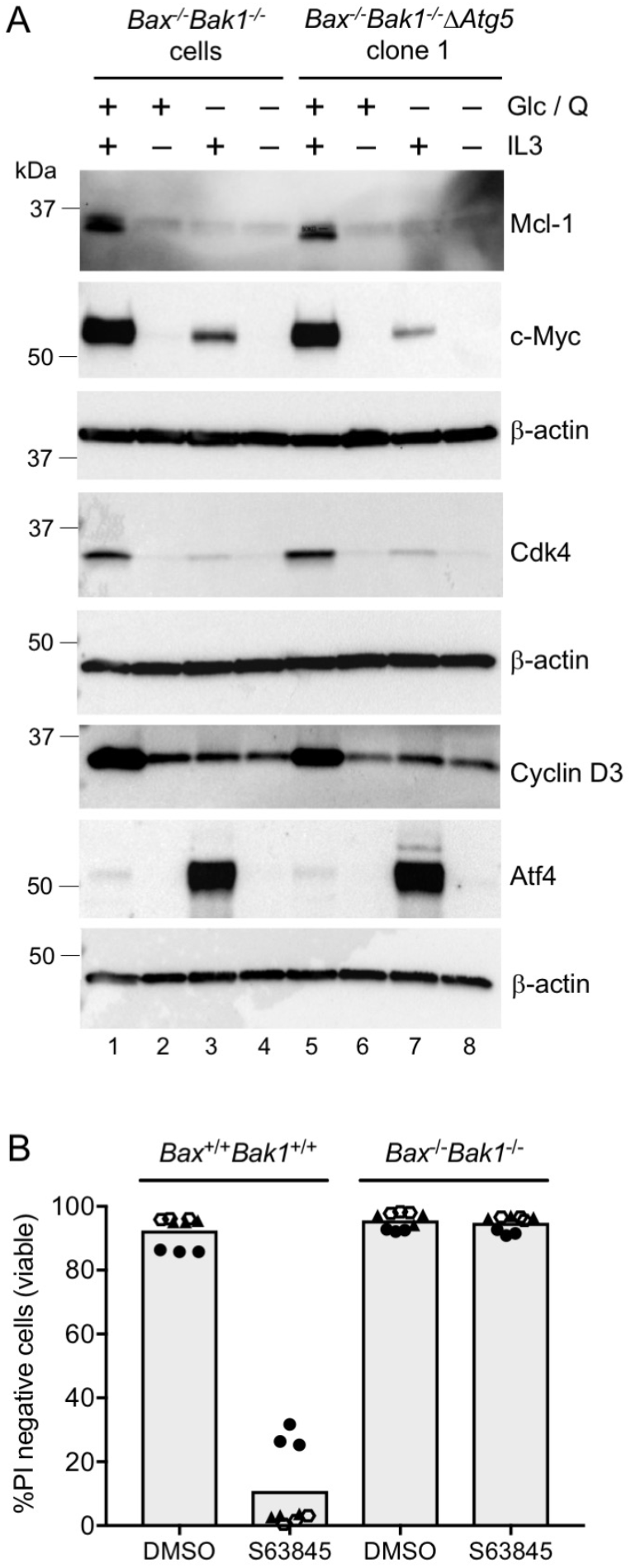
Withdrawal of IL3 or glutamine reduced the expression of Mcl-1 and several key regulators of cell cycle progression. **A.** *Bax*^-/-^*Bak1*^-/-^ cells and *Bax*^-/-^*Bak1*^-/-^Δ*Atg5* clone 1 were cultured for 72 h in medium supplemented with (+) or without (−) added glucose (Glc) and glutamine (Q), as indicated, in the presence or absence of IL3 (+/−). Whole cell lysates were prepared and analyzed by Western blotting to detect the indicated proteins. Three independent membranes were used for immunoblotting and β-actin loading control blots for each are shown. Membrane #1: anti-Mcl-1 and anti-c-Myc; membrane #2: anti-Cdk4; membrane #3: anti-cyclin D3 and anti-Atf4. **B.** *Bax*^+/+^*Bak1*^+/+^ and *Bax*^-/-^*Bak1*^-/-^ cells cultured in complete medium with IL3 were treated with DMSO (vehicle control) or 10 μM of the Mcl-1 inhibitor S63845 for 24 h. Cells were harvested and viability determined by PI exclusion and flow cytometry. Bars depict the mean of 3 independent experiments represented by the 3 different symbols, performed in triplicate. The experiment represented by the solid circles was performed with S63845 synthesized by SYNthesis Med Chem. Experiments represented by the solid triangles and open hexagons were performed with S63845 synthesized by Active Biochem (see Materials and Methods).

As well as causing arrest, IL3 withdrawal reduced c-Myc protein to barely detectable levels in both *Bax*^-/-^*Bak1*^-/-^ and *Bax*^-/-^*Bak1*^-/-^Δ*Atg5* cells (Fig. 6A). Cdk4 and cyclin D3 are key drivers of cell cycle progression in haematopoietic cell lines^12, 13^, and these proteins were also markedly reduced in the IL3-starved, Bax/Bak1-deficient myeloid cells, suggesting a mechanism for their arrest (Figs. 1C and 2D). Withdrawal of glutamine from these cells also caused cell cycle arrest, even in the presence of IL3 (Fig. 4A). Under these conditions, IL3 was able to stimulate the expression of c-Myc and Cdk4, but the levels of these proteins were significantly lower than in the cells cultured with both glutamine and IL3 (Fig. 6A). The transcription factor Atf4 accumulates in response to a variety of cellular stresses including amino acid starvation^14^. In glutamine-starved *Bax*^-/-^*Bak1*^-/-^ and *Bax*^-/-^*Bak1*^-/-^Δ*Atg5* cells, Atf4 levels increase strongly, but intriguingly only in the presence of IL3. These results indicate that the signal transduction pathway activated by ligation of the IL3 receptors remains largely intact in cells cultured without supplemented glutamine, and can stimulate the cells to remain large, but that glutamine is essential for high levels of c-Myc and Cdk4/cyclin D3 expression, and for the cells to divide.

### Bax^-/-^Bak1^-/-^ myeloid cells consume extracellular glucose to maintain long-term viability

The Bax/Bak1-null cells cultured without added glucose and glutamine would need a source of energy to survive (Figs. 4B and S2B), so we tested whether they were obtaining it from the supplemented FBS. In IL3-containing medium lacking added glucose, glutamine and FBS, all *Bax*^-/-^*Bak1*^-/-^ and *Bax*^-/-^*Bak1*^-/-^Δ*Atg5* died after 2 days (Fig. 7A). In the absence of serum, we asked if adding back glutamine or glucose alone was sufficient to support cell survival. While <40% of cells cultured in medium supplemented with 4mM glutamine remained viable after 2 days (and all died within 4 days, Supplementary Fig. S3A), >90% of cells cultured with 5.5 mM glucose survived (for more than 6 days, Supplementary Fig. S3B). In IL3-starved cultures, we observed a similar dependence on glucose, but not glutamine, for cell viability (Fig. 7B). Strikingly, we found that 0.5 mM glucose only (an equivalent amount present in 10% FBS) was sufficient to support the survival of >80% of IL3-deprived *Bax*^-/-^*Bak1*^-/-^ and *Bax*^-/-^*Bak1*^-/-^Δ*Atg5* cells (Fig. 7B) for at least 1 week (Fig. 7C). Collectively, these results suggest that in the absence of growth factor, *Bax*^-/-^*Bak1*^-/-^ myeloid cells consume extracellular glucose to maintain long-term viability, without the need to invoke Atg5-dependent autophagy.

**Figure 7.**
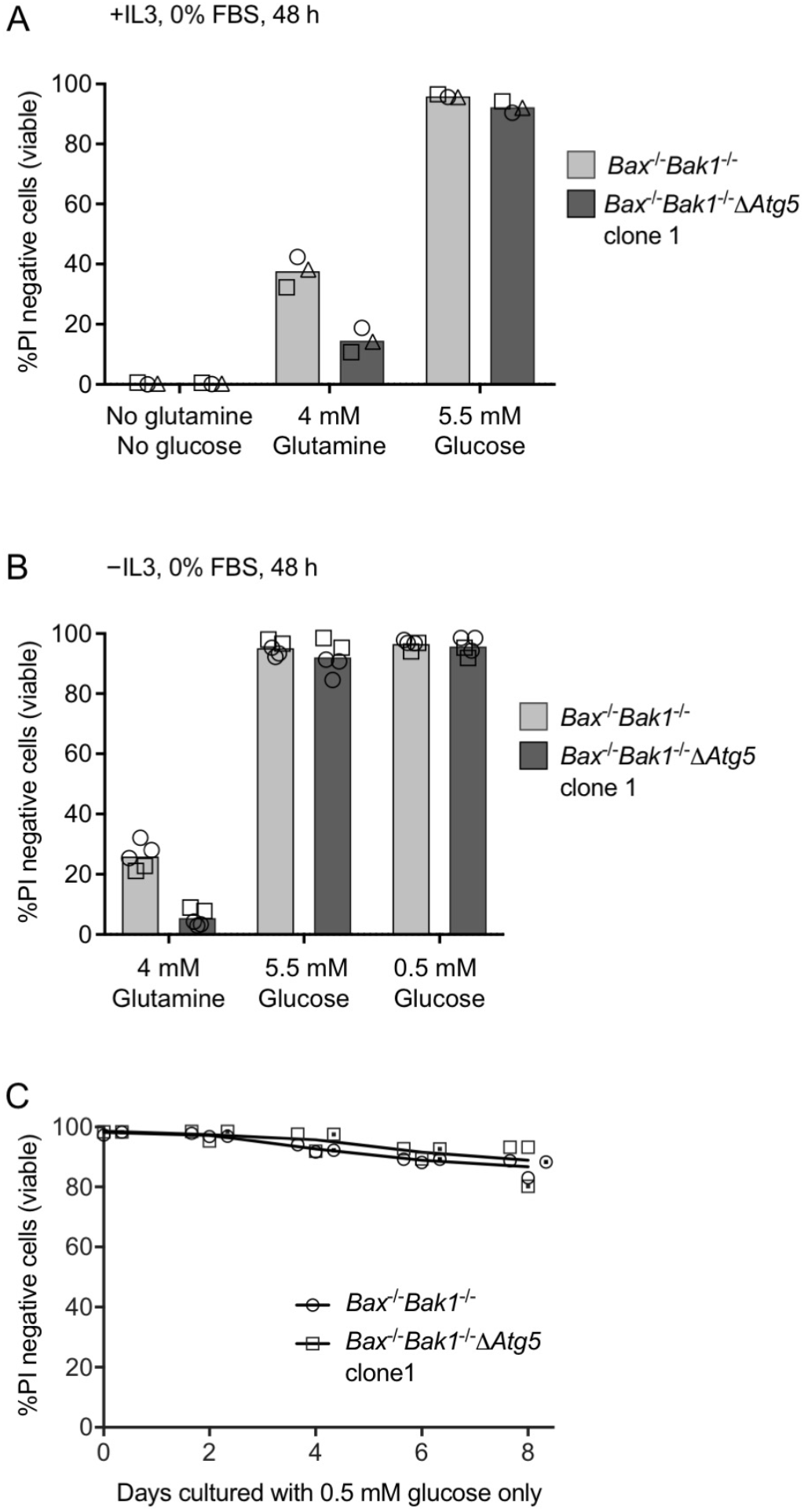
In the absence of Atg5-dependent autophagy, IL3-deprived *Bax*^-/-^*Bak1*^-/-^ myeloid cells consume extracellular glucose to maintain long-term viability. **A.** *Bax*^-/-^*Bak1*^-/-^ cells and *Bax*^-/-^*Bak1*^-/-^Δ*Atg5* clone 1 were cultured in the absence of FBS for 48 h in medium without or with the specified concentration of glutamine and/or glucose, with IL3. Cells were assayed for viability by PI exclusion and flow cytometry. Bars represent the average of 3 independent experiments (depicted by the 3 different symbols). **B.** *Bax*^-/-^*Bak1*^-/-^ cells and *Bax*^-/-^*Bak1*^-/-^Δ*Atg5* clone 1 were cultured in the absence of FBS for 48 h in medium with the specified concentration of glutamine or glucose, without IL3. Cells were assayed for viability by PI exclusion and flow cytometry. Bars represent the average of 2 independent experiments (depicted by the 2 different symbols) performed with technical replicates as indicated. **C.** *Bax*^-/-^*Bak1*^-/-^ and *Bax*^-/-^*Bak1*^-/-^Δ*Atg5* cells were cultured in medium supplemented with 0.5 mM glucose; without IL3, glutamine or FBS. At the indicated times, cells were assayed for viability by PI exclusion and flow cytometry. The curves connect the average values of data points derived from 2 independent experiments (depicted by the 2 different symbols) performed with technical replicates or without (dotted symbols) as indicated.

## Discussion

IL3-dependent myeloid cell lines have provided many insights into the regulation of fundamental processes such as cell death, cell division, and autophagy, as well as the signaling pathways that regulate them. For example, it was experiments on an IL3-dependent myeloid cell line that first revealed that the function of Bcl-2 was to inhibit cell death^3^. When genes for Bax and Bak1 are deleted in factor-dependent myeloid (FDM) cells, they fail to undergo apoptosis when IL3 is removed, but nevertheless will die if starved of nutrients needed to maintain their metabolism. When *Bax*^-/-^*Bak1*^-/-^ FDMs are cultured without IL3, they decrease expression of the glucose transporter Glut1, shrink and arrest, but do not die^4^. This led to the proposal that the cells could not take up glucose, so they needed autophagy to maintain their metabolism to survive. Consistent with this idea, Lum *et al*. found that inhibition of autophagy either by 3-methyladenine, chloroquine or shRNA knockdown of Atg5 caused the cells to die^4^. However, we also used chloroquine to inhibit autophagy and shRNA to reduce Atg5 levels, and found that Atg5-dependent autophagy was not required for the survival *Bax*^-/-^*Bak1*^-/-^ myeloid cells deprived of IL3^15^. It is conceivable that the two Atg5-targeting shRNA sequences used by Lum *et al*., which is different to our shRNA, had off-target effects (such as induction of interferons) that sensitized their cells to autophagy inhibition. Notwithstanding, it is unlikely that off-target effects of chloroquine caused such contrasting survival outcomes in similar IL3-dependent lines.

In this report, we have genetically eliminated *Atg5* expression in IL3-dependent *Bax*^-/-^*Bak1*^-/-^ cells using CRISPR/Cas9. Unlike Lum *et al*., who concluded that autophagy was necessary to sustain the survival of IL3-deprived *Bax*^-/-^*Bak1*^-/-^ FDM cells beyond 4 days^4^, our results demonstrate that canonical Atg5-dependent autophagy pathways are not required to support the long-term viability of these cells (Fig. 2). We generated IL3-dependent FDM lines by co-culturing E14.5 fetal liver single-cell suspensions from *Bax*^-/-^*Bak1*^-/-^ embryos with fibroblasts expressing a *HoxB8* retrovirus (to immortalize the cells) in the presence of IL3^16^. Lum *et al*. obtained IL3-dependent myeloid progenitors from the bone marrow of adult *Bax*^-/-^*Bak1*^-/-^ mice, but did not specify how these cells were immortalized^4^. It is plausible that differences in the age, organ of origin, or immortalization protocols used to generate IL3-dependent myeloid progenitors caused the discrepancy in results between our study and those reported by Lum and colleagues. An experiment to rule out these differences would be to mutate *Bax, Bak1* and *Atg5* genes by CRISPR/Cas9 in IL3-dependent myeloid lines isolated from the fetal livers and bone marrow of wild-type mice, then determine whether these cells survive long-term in the absence of IL3.

We were able to confirm the observation that culture of FDM cells without IL3 reduced the levels of the glucose transporter Glut1^4^, and wondered if the mechanism for reduction of Glut1 would be similar to that described by Wu *et al*. who found that thioredoxin interacting protein (Txnip) could reduce the levels of Glut1 in human hepatocytes^7^. Although we found that Txnip increased in FDM cells when IL3 was removed, levels of Glut1 decreased by the same amount when IL3 was removed from Txnip-null cells, and over-expression of exogenous Txnip did not reduce the levels of Glut1 (Fig. 3). Although we confirmed the reported inverse correlation between levels of Txnip and Glut1, Txnip is neither required nor sufficient for IL3 to regulate levels of Glut1 in FDMs. It remains possible that the Txnip-mediated downregulation of Glut1 is a mechanism specific to hepatocytes and other cell types in response to glucose availability^7^.

Canonical autophagy was shown to be required for the efficient surface expression of Glut1^6^. Our Atg5-deficient *Bax*^-/-^*Bak1*^-/-^ FDM cells had similar levels of Glut1 to parental *Bax*^-/-^*Bak1*^-/-^ cells, and they decreased Glut1 to a similar extent when cultured without IL3. Furthermore, the *Bax*^-/-^*Bak1*^-/-^Δ*Atg5* cells showed similar levels of glucose uptake to *Bax*^-/-^*Bak1*^-/-^ cells when cultured with IL3 and without. Therefore, in FDM cells, Atg5-dependent autophagy is not needed to maintain normal levels of Glut1, and does not affect the ability of IL3-starved cells to take up and metabolize glucose (Figs. 3C and S1D).

Consistent with the requirement of glutamine for lymphoid cell survival^10^, we found that wild-type factor dependent myeloid cells died following glutamine deprivation (Fig. 4). Surprisingly, a high concentration of glutamine was not required to sustain the survival of Bax/Bak1-null FDM cells in serum-containing cultures not supplemented with extra glucose, and that these glutamine-deprived cells maintained a large size in the presence of IL3, but arrested. This suggests that glutamine provided the source of amino acids needed for cell division, but that in its absence, these cells import the glucose present in serum to maintain enough metabolic activity to survive (Fig. 7). Hecht *et al*.^17^ reported that the acute shrinkage of B an T lymphocytes following growth factor withdrawal occurred before the induction of autophagy. Consistent with Hecht *et al*., the shrinkage of FDM cells following growth factor or glutamine withdrawal also occurred independently of Atg5-mediated autophagy (Fig. 5).

In the absence of high levels of glutamine, pathways activated by IL3 could still converge to induce the expression of c-Myc in Bax/Bak1-null FDM cells, and although this was response was blunted, the arrested cells maintained a relatively large size for at least 1 week (Fig. 5). We found that the transcription factor Atf4 was strongly induced by IL3 in glutamine-starved cells, but at present, it is unclear how or why signaling from the IL3 receptor leads to increased Atf4 production. Although Atf4 was shown to activate the expression of several autophagy genes to promote cell survival^18^, we have not been able to confirm that its accumulation induced LC3B lipidation in the glutamine-starved FDM cells. That IL3 also induced Atf4 expression in Atg5-deficient *Bax*^-/-^*Bak1*^-/-^ cells following glutamine deprivation (Fig. 6), suggests that Atf4 does not promote canonical autophagy to maintain cell viability under these conditions. Moreover, cells starved of both IL3 and glutamine showed no detectable expression of Atf4, yet survived for weeks.

Several pathways have been proposed to explain how engagement of IL3 receptors prevents activation of Bax and Bak1. For example, it was proposed that IL3 receptors signaled via Akt to phosphorylate Bad, preventing it from binding to Bcl-x, to thereby free Bcl-x to inhibit apoptosis^19, 20^. However, experiments in which genes for Bad or Akt have been mutated have shown that unlike Bax/Bak1, they play relatively minor roles in the regulation of apoptosis by IL3^16, 21^. These experiments, and those where genes for all the Bcl-2 family member were deleted^22^ suggest that Bax/Bak1 dependent apoptosis is not always signaled via BH3-only proteins, but more often regulated by controlling the abundance of anti-apoptotic Bcl-2 family members, such as Bcl-2, Bcl-x, and Mcl-1^23^. The results presented here are consistent with this, and show that apoptosis of FDMs following the removal of IL3, or culture in the absence of glutamine, is due to the decrease in abundance of Mcl-1. Cytokines including IL3 keeps cells alive by driving *Mcl-1* transcription, most likely via Jak2 and Stat5^24, 25, 26^, and glutamine is required to maintain the production of Mcl-1 to prevent default apoptosis. Consistent with a requirement for Mcl-1 to prevent spontaneous activation of Bax/Bak1, a specific inhibitor of Mcl-1 was able to induce apoptosis of wild-type *Bax*^+/+^*Bak1*^+/+^ FDM cells cultured with IL3 and glutamine (Fig. 6).

In the absence of growth factor and high concentrations of added glucose and glutamine, Atg5-deficient *Bax*^-/-^*Bak1*^-/-^ cells remained viable for weeks in serum-containing medium, and their rapid death in glucose-free cultures was consistent with the hypothesis that they took up and used the glucose present in serum to survive (Fig. 7). That these cells predominantly generate chemical energy through glycolysis (Fig. 3) was also consistent with their ability to consume only diluted concentrations of glucose to remain viable for at least a week. Serum-starved Bax/Bak1-null cells supplemented with high concentrations of glucose and glutamine survived for several weeks, but arrested, with or without IL3 (data not shown). Collectively, our work suggests that in the absence of growth factor, *Bax*^-/-^*Bak1*^-/-^ FDM cells, including mutant lines lacking Atg5-dependent autophagy, import extracellular glucose and amino acids to maintain long-term viability. Their metabolic requirement for glucose and amino acids is a mechanism to support survival even in cultures where levels of these nutrients are not optimal for cell proliferation, and the reduced levels of c-Myc, Cdk4 and cyclin D3, combined with possibly altered levels of other cell cycle regulators, could explain why growth factor and/or nutrient-starved *Bax*^-/-^*Bak1*^-/-^ cells arrested (Fig. 8)

**Figure 8.**
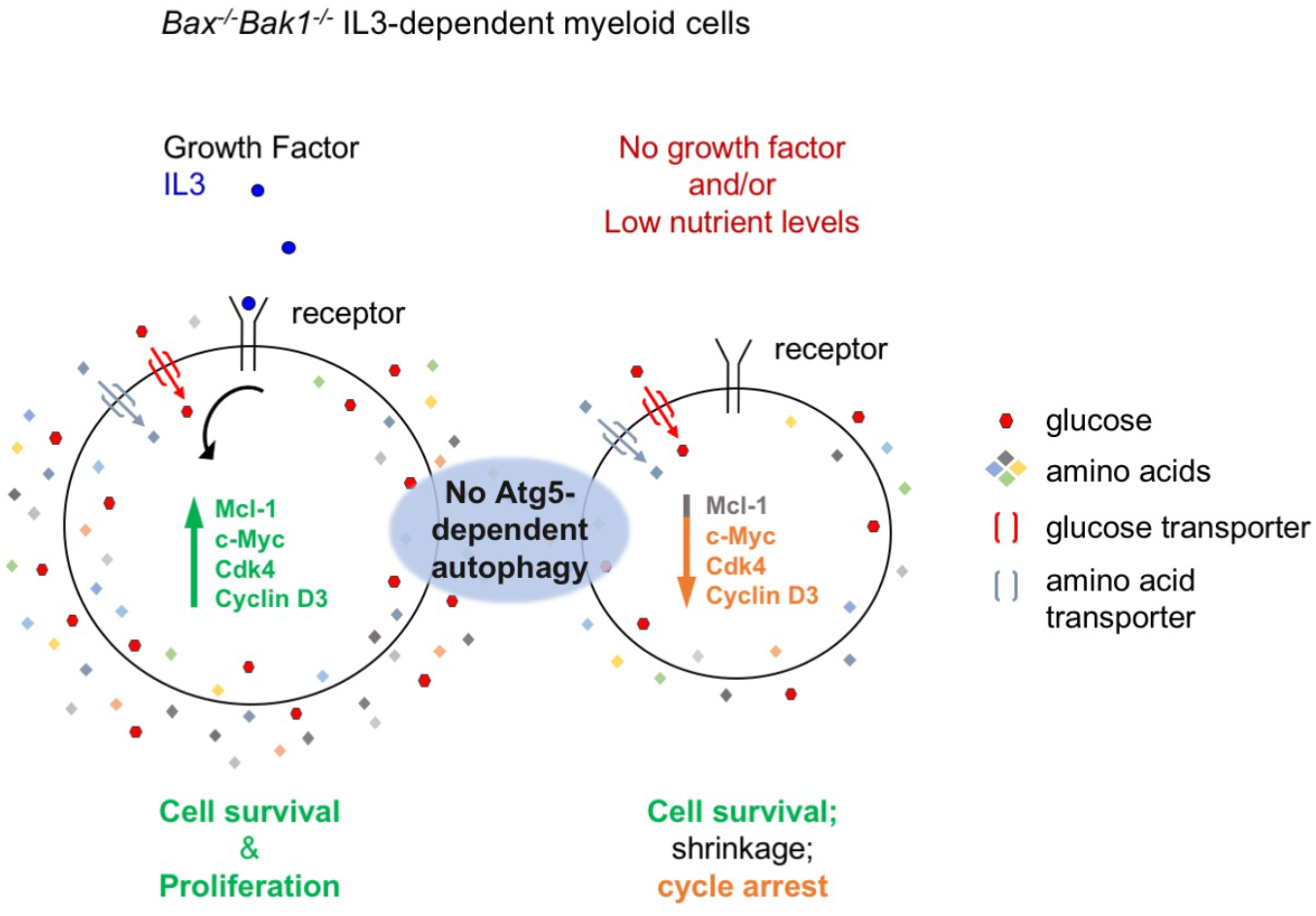
The growth factor IL3 stimulates the expression of Mcl-1 that inhibits activation of Bax/Bak1, and stimulates expression of c-Myc, Cdk4 and cyclin D3 to promote cell growth and division. Optimal concentrations of glucose and glutamine are required to maintain levels of these proteins needed to prevent activation of Bax/Bak1 (Mcl-1), as well as for cell proliferation (c-Myc, Cdk4, cyclin D3 and possibly others). In the absence of growth factor, or when nutrient levels are low, sufficient levels of glucose transporters (Glut1, Glut4 and possibly others) remain to allow *Bax*^-/-^*Bak1*^-/-^ myeloid cells to consume extracellular glucose to survive, even in cells lacking Atg5-dependent autophagy.

## Methods & Materials

### Cell culture

Wild-type *Bax*^+/+^*Bak1*^+/+^ and *Bax*^-/-^*Bak1*^-/-^ IL3-dependent mouse myeloid progenitor (FDM) cell lines have been described^16^. We routinely test our cells for mycoplasma contamination. Cells were cultured in Dulbecco’s Modified Eagle Medium (DMEM, ThermoFisher Scientific Cat# 11885-084; containing 5.55 mM D-glucose and 4 mM L-glutamine) supplemented with 10% fetal bovine serum (FBS, Sigma Cat# F9423) and 0.25 ng/mL recombinant murine interleukin 3 (IL3, PeproTech, Cat# 213-13) in a 37°C humidified incubator with 10% CO2. Cells were deprived of IL3, glucose, glucose and glutamine, or varying concentrations of FBS, by washing 3 times with medium that lacked those nutrients, then resuspended in the same medium for analysis at the indicated time points. Unless indicated otherwise, 1 x 10^5^ cells were seeded in 1 mL of the specified medium per well of a 24-well tissue culture plate (Corning, Cat# 353047), and one well was analysed for each timepoint. For experiments without FBS, plastic tubes and tissue culture dishes were pre-coated with 20 μg/mL bovine serum albumin (Sigma Cat# A1470) and rinsed with serum-free medium before cell seeding. DMEM lacking D-Glucose and sodium pyruvate but containing 4 mM L-Glutamine (Cat# 11966-025) and DMEM lacking D-Glucose, sodium pyruvate, L-Glutamine and phenol red (Cat# A14430-01) were purchased from ThermoFisher Scientific. D-Glucose was purchased from Sigma (Cat# G7528). The glucose concentration in FBS was determined using an Accu-Chek Performa glucose meter, according to the manufacturer’s instructions.

### Lentiviral constructs for CRISPR/Cas9 and inducible overexpression

An inducible lentiviral CRISPR/Cas9 system^27^ was used to mutate *Txnip* in *Bax*^-/-^*Bak1*^-/-^ FDM cells. Guide RNA oligos targeting mouse *Txnip* (guide 1: CCA GCA GGT GAG AAC GAG A; guide 2: TAT GGT TGC GTA GAC TAC T) were cloned into the *Bsm*BI site of the pFgH1tUTG GFP lentiviral vector. Guide RNA oligo targeting mouse *Atg5* (AAG ATG TGC TTC GAG ATG TG) was cloned into the *Bsm*BI site of the lentiviral vector lentiCRISPR v2 (Addgene #52961^28^). For overexpression studies, the coding sequence of mouse *Txnip* was PCR amplified from pEGFP-C1-Txnip (Addgene #18758^29^) with the incorporation of an N-terminal FLAG-tag. The PCR product was cloned into the £coRIIΛ%eI sites of a doxycycline inducible pFTREtight MCS rtTAadvanced GFP lentiviral vector^21^.

### Antibodies and small molecules

Antibodies: LC3B (D11, Cell Signaling Technology #3868), Atg5 (D5F5U, Cell Signaling Technology #12994); β-actin (AC-15, Sigma #A1978), Glut1 (Abcam ab15730), Txnip (D5F3E, Cell Signaling Technology #14715S), Glut4 (1F8, Cell Signaling Technology #2213), Mcl-1 (Rockland 600-401-394S), c-Myc (Abcam ab32072), Cdk4 (Santa Cruz Biotechnology sc-260), Cyclin D3 (Cell Signaling Technology #2936), Atf4 (D4B8, Cell Signaling Technology #11815). Small molecules: Mcl-1 inhibitor S63845 was synthesized according to Kotschy *et al*,^11^ by SYNthesis Med Chem Pty Ltd (Parkville, Australia) or by Active Biochem (Catalog# A-6044 – a kind gift from David Huang); DMSO (dimethyl sulfoxide; Cat# 472301) and propidium iodide (Cat# P4864) were purchased from Sigma.

### Measurements of cell death, cell cycle and cell size

Cell death and cell cycle were assayed using propidium iodide (PI) and flow cytometry. For death assays, cells were harvested and resuspended in PBS containing 1 μg/mL PI. For cell cycle analysis, cells were harvested and resuspended in PBS containing 0.1% Triton X-100, 5 μg/mL PI and 10 μg/mL RNaseA (Thermo Scientific), then incubated at 37 °C for 15 min. Samples were processed on FACSCalibur or LSRFortessa X-20 flow cytometers (BD Biosciences) and the acquired data analyzed with FlowJo software. Cell volume in Figure 5 was measured on a CASY instrument (Roche Innovatis AG), a Coulter-based cell counter operating on the Electrical Current Exclusion principle. In brief, 200 μL aliquots of cells cultured under the specified conditions were mixed with 5 mL of CASY electrolyte buffer and analyzed in a 60 μm CASY capillary. Multiple parameters including the peak cell volume were collected.

### Glucose uptake assays

Extracellular acidification (ECAR) rates were measured in live cells using a Seahorse Bioscience XFe96 Analyzer (Agilent Technologies). In brief, cells were resuspended in non-buffered DMEM (Seahorse Biosciences) and 200 x 10^3^ cells were seeded per well in Seahorse Bioscience 96-well plates. The cellular ECAR was analyzed in non-buffered DMEM containing 2 mM glutamine with the following compounds: 5 mM glucose, 1 μM oligomycin and 50 mM 2-deoxy-glucose (2DG). For each assay cycle, measurement time points of 2 min mix, 2 min wait and 5 min measure were collected.

### Data reproducibility

Unless indicated otherwise, all experiments shown have been reproduced at least 3 independent times with similar results. While not all repeats were performed wholly or in exact detail, we ensured that our experimental design allowed us to interpret and draw conclusions from multiple independent experiments. This is aided by the fact that our results are unambiguous: the majority of cells are either alive or dead, cycling or arrested, large or small; and immunoblots show clear changes in protein levels or modification. We have endeavored to perform experiments that provide consistent evidence in mechanistically independent ways, rather than repeating the same experiment many times until statistical significance is reached. The experimental triangulation of our evidence^30^ is best exemplified by multiple independent assays confirming that autophagy-deficient FDM cells survived for weeks without growth factor under a variety of different culture conditions (Figs. 2B, 4B, 5, 7C, S3B).

## Acknowledgments

We thank Gabriela Brumatti, Rebecca Feltham, James Vince, Simon Monard, David Huang and Andre Samson for discussion, technical advice and reagents. Funding for this project was provided by the Australian National Health and Medical Research Council (NHMRC) Program Grant #1113133 and NHMRC fellowship 1020136 to DLV; and the Leukemia and Lymphoma Society SCOR grant #7001-13. Work in the authors’ laboratory is made possible by operational infrastructure grants through the Australian Government Independent Research Institutes Infrastructure Support (IRISS) and the Victorian State Government OIS.

